# Maximising the translational potential of neurophysiology in amyotrophic lateral sclerosis: a study on compound muscle action potentials

**DOI:** 10.1101/2024.05.09.593349

**Authors:** Scott McKinnon, Zekai Qiang, Amy Keerie, Tyler Wells, Pamela J. Shaw, James J.P. Alix, Richard J. Mead

**Affiliations:** Department of Neuroscience, Sheffield Institute for Translational Neuroscience (SITraN), The University of Sheffield, Sheffield, S10 2HQ; Neuroscience Institute, The University of Sheffield, Western Bank, Sheffield, S10 2TN

## Abstract

Transgenic mouse models of amyotrophic lateral sclerosis, such as the widely used SOD1^G93A^ mouse, enable investigation of disease mechanisms and testing of novel therapeutic interventions. However, treatments that have been considered successful in mice have often failed to translate into human benefit in clinical trials, particularly when relying on the so-called ‘survival’ read-out. Compound muscle action potentials (CMAPs), are a simple neurophysiological test that measures the summation of muscle fibre depolarisation in response to maximal stimulation of the innervating nerve. CMAPs can be measured in both mice and humans and decline with motor axon loss in ALS, making them a potential translational read-out of disease progression which could help bridge the preclinical and clinical divide. Herein we assess the translational potential of CMAPs and ascertain at what time points human and mouse data aligned most closely. We extracted data from 18 human studies and compared with results generated from SOD1^G93A^ and control mice at different ages across different muscles. We found that the relative CMAP amplitude difference between SOD1^G93A^ and control mice in tibialis anterior and gastrocnemius muscles at 70 days of age was most similar to the relative difference between baseline ALS patient CMAP measurements and healthy controls in the abductor pollicis brevis (APB) muscle. We also found that the relative decline in SOD1^G93A^ tibialis anterior CMAP amplitude between 70-140 days was similar to that observed in 12 month human longitudinal studies in APB. Our findings suggest CMAP amplitudes can provide a ‘translational window’, from which to make comparisons between the SOD1^G93A^ model and human ALS patients. CMAPs are easy to perform and can help determine the most clinically relevant starting/end points for preclinical studies and provide a basis for predicting potential clinical effect sizes.

## Introduction

In amyotrophic lateral sclerosis (ALS) an increasing number of molecules are being taken from preclinical studies into clinical trials. While many therapeutic approaches have been successful in mice but unsuccessful in humans^1–6^, both edavarone and the antisense oligonucleotide Tofersen have transitioned into positive human trials from successful preclinical studies^7,8^. While modelling the human disease in mice poses many problems^9^, several models are available which provide a testing platform for different pathophysiological aspects of the disease. These include the ALS-FTD TDP-43^Q331K^ knock-in^10^ and transgenic models^11^, humanised FUS models^12^ and the most widely studied model, the SOD1^G93A^ mouse, which provides a robust, predominantly lower motor neuron phenotype^13,14^. Bacterial artificial chromosome (BAC) transgenic models carrying an ALS C9orf72 allele with a hexanucleotide repeat expansion have shown phenotypes in some laboratories^15^ but not others^16^.

Despite obvious advantages, relatively few therapeutic candidates have been studied with the same read-out of disease in both preclinical and clinical trials. In the SOD1 model, time to reach end-stage disease (“survival time”) is often used^1,3,6^, partly because this is a common regulatory requirement in Phase III clinical trials^17^. However, positive effects on survival can be the result of poor experimental design^18^. Evaluation of other outcome measures could provide enhanced insight into treatment effects and may provide a better platform for assessing the potential for positive clinical outcomes. For example, in clinical trials, the most common measure of disease state is the amyotrophic lateral sclerosis functional rating scale-revised (ALSFRS-R)^19^. However, while there are measures of symptom severity in mice (e.g. neurological score^14^), these are not directly translatable.

Neurophysiological tools are attractive candidates for translational studies, as the same measurements can be made in both mice and humans. Compound muscle action potential (CMAP) amplitude is one example, representing the balance between motor unit loss and compensatory reinnervation. As a result, CMAP amplitude is less sensitive to motor neuron loss compared to motor unit estimation measures^20–24^. However, it is arguably the simplest neurophysiological readout to incorporate into complex preclinical studies.

To explore the potential of neurophysiology to provide a translational biomarker of disease, we have compared CMAP amplitude data collected from a systematic search of the available literature to our own CMAP data from multiple muscles in the SOD1^G93A^ mouse. Our first aim was to assess which muscles and timepoints in the mice align best with ALS patient CMAP data at the time of recruitment into clinical studies. Second, we sought to establish which muscles in SOD1^G93A^ mice demonstrate a decline in CMAP over time that is most similar to that observed in ALS patients.

## Methods

### Ethics

All *in vivo* studies were carried out in compliance with the Animals (Scientific Procedures) Act 1986 and all work was completed by UK Home Office personal licence holders under a UK Home Office project licence. All mice were housed and cared for according to the Home Office Code of Practice for the Housing and Care of Animals Used in Scientific Procedures.

### Mouse Model

Transgenic SOD1^G93A^ C57BL/6 mice have been described previously^14^. Mice contain ∼23 copies of the human transgene containing the Glycine to Alanine substitution at Amino Acid 93 as described by Gurney et al.^13^.

Mice were housed in groups of 2 to 5 mice with ad libitum access to food (Envigo) and water. The temperature was maintained at 21°C with a 12-hour light/dark cycle.

### Genotyping

Mice were identified by ear clipping and genomic extraction using QuickExtract (Lucigen) and incubating at 65°C for 15 minutes and 98°C for 2 minutes. Genotyping PCR was performed on extracted DNA in 10μl mastermix containing: 2μl FirePol (Solis BioDyne); 0.25μl of 10μM human SOD1 primers (forward 5’-CATCAGCCCTAATCCATCTGA-3’, reverse 5’-CGCGACTAACAATCAAAGTGA-3’); 1μl of 10μM control IL2 primers (forward 5’-CTAGGCCACAGAATTGAAAGATC-3’, reverse 5’-GTAGGTGGAAATTCTAGCATCATCC-3’); 5μl of nuclease-free water; and 0.5μl of template DNA. Following PCR, samples were run on a 2% agarose gel. IL2 and human SOD1 PCR products were visualised at ∼320bp and ∼250bp, respectively.

### Electrophysiological Recording

Electrophysiological testing was carried out at P70, P105 and P140 in 15 transgenic SOD1^G93A^ C57BL/6 (T) and 15 control C57BL/6 female mice (NT). Mice were anaesthetised using gaseous isoflurane (5% isoflurane, flow rate 4L/min oxygen) and then maintained under gaseous anaesthesia (1-2% isoflurane, 0.5L/min oxygen continuous inhalation through a nose cone for the duration of the experiment). Recording electrodes (Ambu Neuroline) were placed in either the tibialis anterior, gastrocnemius, intrinsic muscle of the foot, or the biceps muscles. A reference electrode was placed into the Achilles tendon area for hind limb recordings, and the wrist for forelimb recordings. A ground electrode was placed in the base of the tail. A pair of stimulating electrodes were placed on the skin above the sciatic nerve for hindlimb recordings and above the musculocutaneous nerve for forelimb recordings. Recordings were made using a Dantec Keypoint Focus EMG system. A 0.1ms square pulse was used and the current gradually increased to obtain supramaximal compound muscle action potential (CMAP) responses. The CMAP amplitude was used to measure the output of the targeted muscles. Data were calculated relative to control non-transgenic (NTg) mice at each timepoint and as relative decline from 10 weeks of age. GraphPad Prism version 9.3.1 was used for all preclinical statistical analysis (GraphPad Software, San Diego, California USA).

### Clinical studies: search strategy

Clinical studies were collated through systematic interrogation of the Medline and EMBASE databases for papers up to June 2022. Additional papers were identified through screening the reference list of returned review articles. The studies were restricted to English language only. The search strategy involved a combination of MeSH Terms and keywords: “motor unit number estimate” OR “compound muscle action potential” OR “motor unit number index” AND “motor neuron disease” OR “amyotrophic lateral sclerosis”. We first searched for papers reporting both patient and healthy volunteer values in the same study. Next, we sought papers reporting patient data longitudinally for 12 months.

### Study selection

The full texts across both searches were included for data extraction and synthesis if they reported mean and standard deviation of human CMAP data in individual limb muscles. For the comparison of healthy volunteers and ALS patients, the presence of a healthy control group at a single time point was required. Studies in which ALS patients were not age and gender matched with healthy controls were excluded. For longitudinal data studies, CMAP values were required up to and including 12 months^25–30^.

### Data extraction

For each included study, data regarding author, publication year, muscles studied, healthy control inclusion criteria, matching status, diagnostic criteria, ALSFRS-R score, and subgroup stratification were extracted. Sample size, mean with standard differences for age, CMAP amplitudes of individual muscles and disease duration were recorded for both healthy and ALS cohorts.

### Synthesis

For single studies, extracted raw means and standard deviation data for each muscle were imported. A random-effects model was deployed for quantitative synthesis to take into account the inter-study heterogeneity from varied subject inclusion criteria and demographics. Pooled means were calculated and 95% confidence intervals were determined using Hartung-Knapp adjustment. Heterogeneity between studies was tested by using Higgins and Thompson’s I2 Statistic. For longitudinal studies, percentage changes with respective standard error of the mean (SEM) in CMAP values from baseline were calculated using the extracted data at each timepoint. A linear mixed effects model with a fixed factor “Time (months)”, random factor “Study”, and dependent variable “Percentage change from baseline” was established in order to assess disease progression over the course of 12 months^31,32^. These analyses were performed using the statistical programme RStudio2022.12.0+353 (Posit team, Boston, MA USA).

## Results

### Human clinical study selection

A total of 54 unique single time point studies were identified, of which 36 were excluded and 18 included. Reasons for exclusion included: no reported relevant CMAP data (n = 17); unmatched for age and/or gender (n = 12); individual muscle CMAP not reported and/or not specified (n = 3); mean CMAP and standard deviation not reported for individual muscles (n = 3); bulbar muscle recording (n = 1). CMAP measurements were most frequently reported in abductor pollicis brevis (APB; 15 studies). Abductor digiti minimi (ADM) CMAP amplitude was reported in 11 studies, tibialis anterior (TA) in 3, biceps brachii (BB), abductor hallucis (AH), extensor digitorum brevis (EDB) and flexor carpi ulnaris (FCU) in 1 study. The study characteristics are shown in supplemental table 1. There was significant inter-study heterogeneity with I^2^ ≥ 75% across all cohorts, an example of this is shown for APB in supplementary figure 1. In addition, 22 full texts were reviewed for longitudinal CMAP amplitude data at various monthly intervals. Eight studies were excluded, mostly due to a lack of relevant electrophysiological data (n = 6), one did not report CMAP data for individual muscles. APB and ADM were the most frequent muscles to be assessed longitudinally (seven and ten studies respectively).

**Table 1:** Heterogeneity across human studies. *The study divided ALS patients into two cohorts: one with normal CMAP amplitude, the other with reduced CMAP amplitude. **In this study, ALS patients were divided by upper limb onset ALS and flail arm syndrome. ***ALS patients were divided based on the presence of weakness and wasting of intrinsic hand muscles.

| Author | Year | Mean age of healthy controls | Diagnostic criteria | Mean age of ALS patients | Mean disease duration (months) | Mean ALSFRS-R score | Reference Number |
| --- | --- | --- | --- | --- | --- | --- | --- |
| Wang* | 2019 | 51.8±9.6 | Awaji-Shima | 54±13<br>57±15 | 10±8<br>12±12 | 43±5<br>41±6 | 1 |
| Paparella | 2021 | 64.38±9.66 | revised El Escorial | 66.5±8.74 | 28.57±19.77 | 39.28±5.2<br>2 | 2 |
| Escorcio-Bezerra<sup>3</sup> | 2016 | 62.6±7.4 | revised El Escorial | 62.2±8.5 | 19.28±11.53 | NA | 3 |
| Kim* | 2016 | 59.3±8.6 | revised El Escorial | 61±11.5<br>58.9±11.2 | 7.8±7.4<br>10.5±8.3 | 38.9±3.9<br>36.3±3.3 | 4 |
| Zheng | 2019 | 49.3±9.3 | Awaji-Shima | 59.1±12.8 | 16±5.8 | NA | 5 |
| Shibuya | 2016 | 59.9±1.3 | revised El Escorial | 61.5±0.8 | NA | 41.06±0.4 | 6 |
| Singh | 2017 | 25.29±2.3 | revised El Escorial | 51.86±9.55 | 11.14±2.85 | NA | 7 |
| Sharma | 1996 | 48±4 | revised El Escorial | 55±4 | 28±5 | NA | 8 |
| Min | 2020 | 62.98±8.17 | revised El Escorial | 61.57±10.74 | 19.2±19.73 | 36.47±6.2<br>5 | 9 |
| Jenkins | 2018 | 54.2±15.7 | revised El Escorial | 57.1±13.5 | 16.5 (Range 6.25–66.25) | 40±4.5 | 10 |
| Sun** | 2016 | 55 (Range 45–70) | revised El Escorial | 55 (Range 41–76)<br>58 (Range 50–71) | NA<br>NA | NA<br>NA | 11 |
| Oguz-Akarsu<sup>12</sup> | 2022 | 44.9±13 | Awaji-Shima | 53±12.1 | 13.5±13 | 38.7±7.2 | 12 |
| Iwai* | 2016 | 64.2±1.4 | revised El Escorial | 64.6±1.2<br>67.8±1.1 | 13.8±1.2<br>18.1±1.7 | NA<br>NA | 13 |
| Fang*** | 2016 | 50.9±10.86 | revised El Escorial | 47.15±9.56<br>49.6±8.93 | 14.3±6.59<br>15.1±15.77 | 38.2±6.65<br>40.3±3.33 | 14 |
| Fang | 2016 | 50.9±12.8 | revised El Escorial | 52.8±11.4 | 12±9.6 | NA | 15 |
| Deilami | 2019 | 38±11 | revised El Escorial and Awaji-Shima | 39.2±14 | 53.16±13.2 | NA | 16 |
| Gunes | 2021 | 55.8±12.3 | Awaji-Shima | 56.2±12.1 | 9.52±7.6 | NA | 17 |
| Shimizu | 2018 | 65.1±12.3 | revised El Escorial and Awaji-Shima | 66.5±9.3 | 19.2±16.8 | 39.4±7.5 | 18 |
| Boekestein | 2012 | Median 62 (Range 49–78) | revised El Escorial | Median 64 (Range 21–77) | Median 20.4 (Range 9.6–66) | NA | 19 |

### Finding a starting point in mouse to match human baseline data

Using CMAP amplitude data, we first assessed which muscles in the SOD1^G93A^ mouse model, and at which time points, are most representative of the starting point in human clinical trials. We first compared CMAP data from human papers containing both patients with ALS and healthy volunteers (Table 1). While there was significant heterogeneity across studies, CMAP amplitudes in patients were always reduced relative to healthy controls, with APB demonstrating the biggest relative difference (supplementary Fig 1: ALS: 5.6 +/-0.62mV, healthy controls: 10.64 +/-0.74mV; percentage decline of 47.31%).

We then examined the differences between SOD1^G93A^ and healthy mice across different muscles and at different ages and compared those results to the human data (Fig 1). This showed that tibialis anterior and gastrocnemius CMAP amplitudes at P70 in the SOD1^G93A^ mice demonstrated a similar relative difference to ALS patients/healthy controls for the APB muscle. The relative reduction in mouse intrinsic foot muscle CMAP amplitude at P105 was similar to APB and FDI CMAPs in humans. In contrast, biceps brachii CMAPs in the mouse did not map well to human data at any of the ages studied.

**Figure 1.**
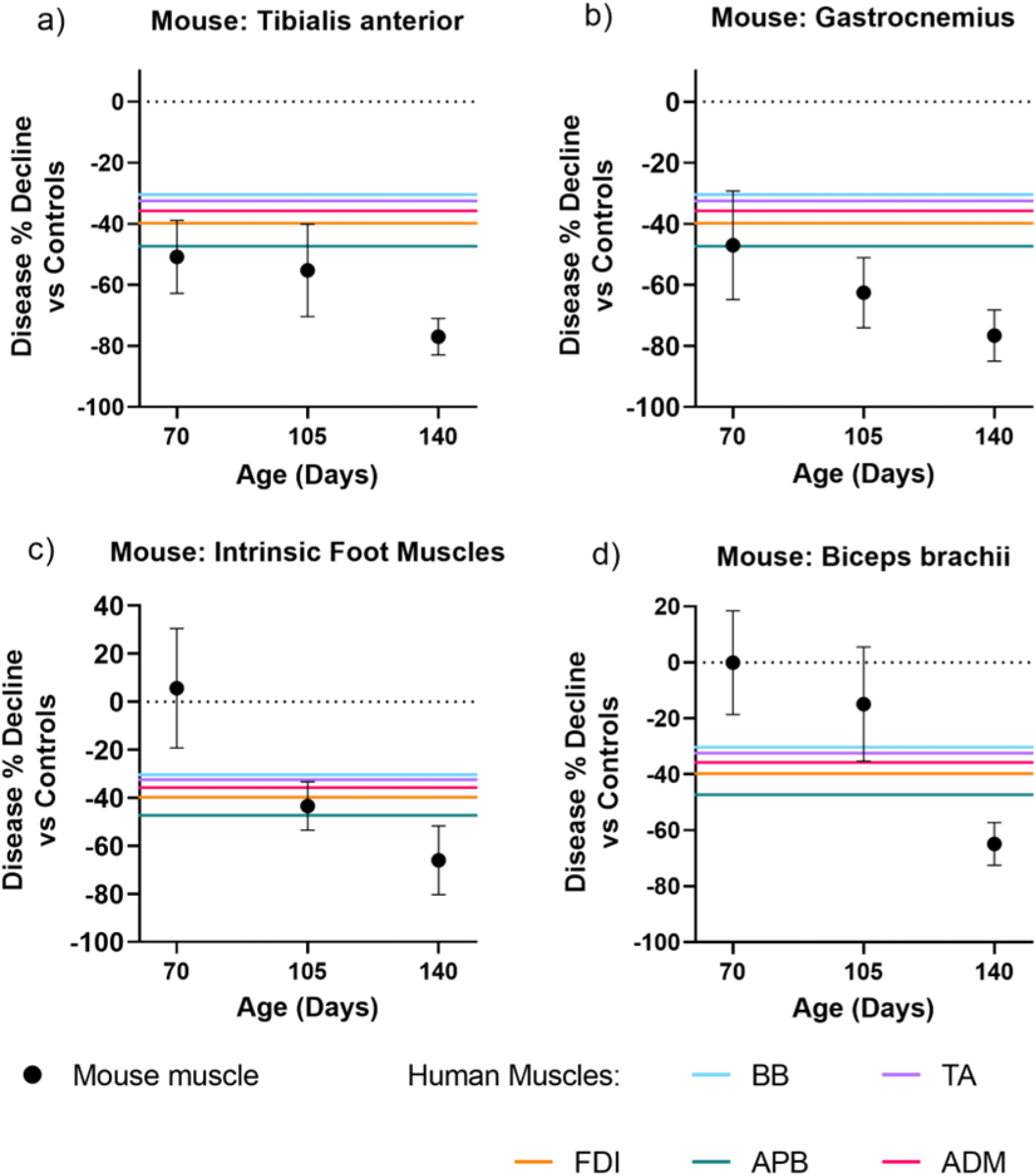
SOD1^G93A^ 70 day tibialis anterior and gastrocnemius CMAP % decline versus healthy controls most resemble human ALS patient versus healthy controls at baseline visit. SOD1^G93A^ mouse data (black circles) is presented as the percentage difference to age-matched non-transgenic controls at three time points (average ± SEM). Similarly, human data is also presented as the relative difference when compared to age/gender matched healthy volunteers. Data are shown for each mouse muscle (black) studied: tibialis anterior (a), gastrocnemius (b), intrinsic foot muscles (c) and biceps brachii (d), overlaid with patient data for different muscles (coloured lines).

### How do CMAP amplitudes change over time in mice and humans?

As expected, CMAP amplitudes declined over time in both human and SOD1^G93A^ mice (Fig 2 and Supplemental Fig 2-6, Supplemental Tables 2 and 3). As the mouse data for tibialis anterior and gastrocnemius at P70 appeared to match the human baseline data in APB, we first compared the relative decline in SOD1^G93A^ and human patients for these muscles (Fig 2). From this analysis it can be seen that the P140 data in mice matches the 12 month time point in human patients and thus the time frame P70-P140 in the mouse would appear to map well to 12 month human studies. Similar plots for the other muscles are given in the supplementary data (Supplementary Fig 2-5). Of these, SOD1^G93A^ tibialis anterior also aligned well to human tibialis anterior (Supplemental Fig 5).

**Figure 2.**
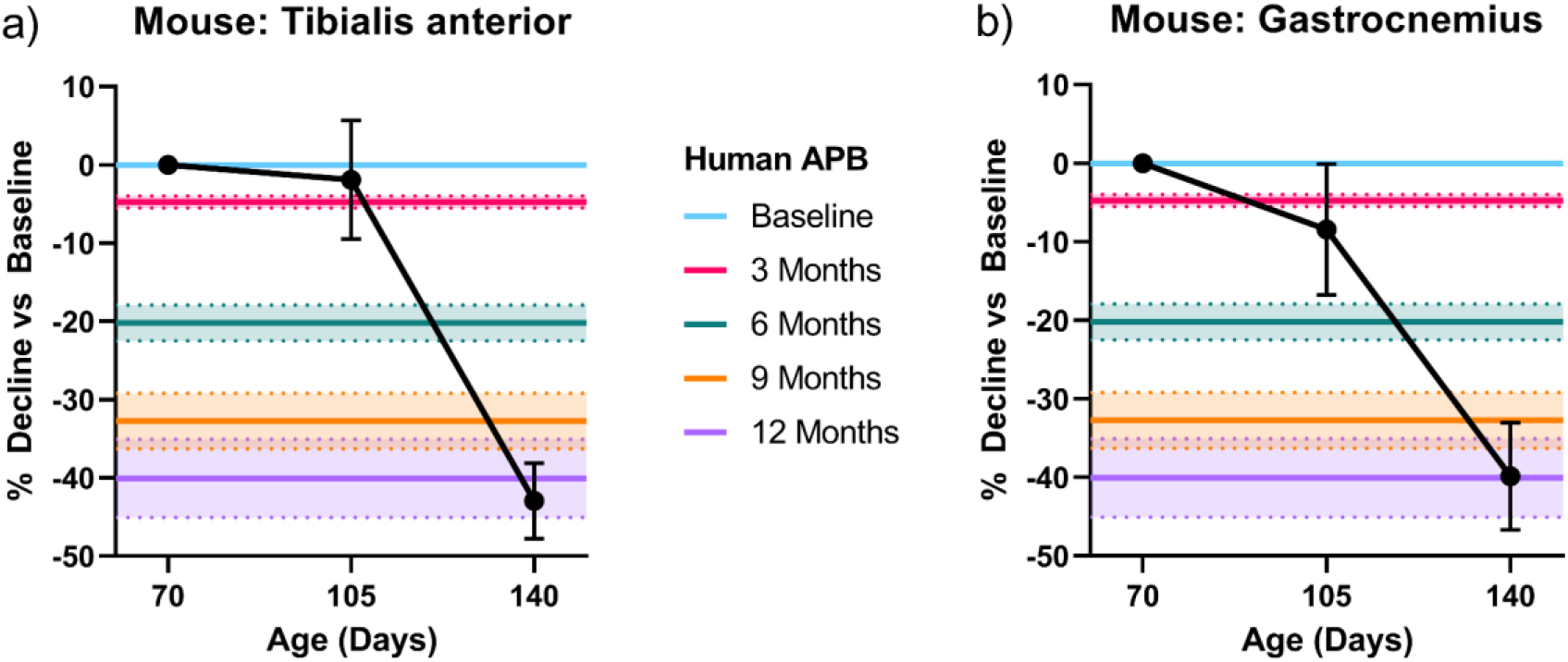
SOD1^G93A^ tibialis anterior and gastrocnemius longitudinal data from P70-P140 represent a good match to the 12 month decline in human APB. Average (± SEM) CMAP decline compared to baseline recording at three time points in the mouse (black circles) in tibialis anterior (a) and gastrocnemius (b). The different time points in human APB are represented by horizontal coloured lines (mean represented by the line and standard error by the shaded area.

## Discussion

In this study we have attempted to align mouse CMAP data with patient data, in order to optimise the translational potential of preclinical CMAP data. Our results suggest that there are some muscles and time points in the mice that are a better representation of human data than others. While there are limitations to our approach, these observations can stimulate debate on how best to measure ALS disease progression in preclinical studies.

Despite the difficulties in translating successful preclinical studies into successful clinical trials, animal models, and in particular mouse models, remain an important element of ALS research. Between 2019 and 2022, around 400 papers per year were published using mouse models of ALS (PubMed search: ((mouse models) OR (mouse model)) AND ((motor neuron disease) OR (amyotrophic lateral sclerosis)). While there are a number of challenges facing drug development, clinical relevance of the data generated in preclinical models has been identified as being of key importance^33^.

There has been a rapid increase in the development and validation of biomarkers for ALS in patients^34,35^. However, the development of preclinical biomarkers has lagged behind the human data, although we note that some clinical biomarkers such as neurofilament levels underwent early preclinical evaluation^36,37^. Neurophysiological measures are one of a number of different approaches that can be applied in both mice and patients. The most obvious example is the extension of the CMAP measurement termed motor unit number estimation (MUNE). There are now several MUNE techniques available for patients and, in mice, older methods such as multi-point and incremental MUNE can be applied^38,39^. We acknowledge that MUNE methods could provide a more sensitive readout of disease state than CMAP amplitude in both mouse models and humans and thus may be preferable when possible. However, as a simple starting point, CMAPs are much easier to perform and can be learnt quickly by scientists whose expertise often lies in other disease-relevant areas.

Our work also has methodological limitations. For example, we have included multi-centre human data, but only single centre mouse data. While our CMAP declines are similar to those reported elsewhere^39^, the addition of preclinical data from other laboratories would strengthen the analysis. Furthermore, we have recorded CMAPs in mice using needle electrodes, while surface electrodes are used in human subjects. Comparing differences relative to healthy controls, or relative to a defined starting point, in both mice and patients helps offset this difference. We also saw significant heterogeneity in human data which highlights the importance of having well defined recording protocols in multi-centre studies. In human studies there is also the added complexity of recruiting patients at different stages of disease and with different disease trajectories. Recent modelling work has demonstrated several different patterns of ALSFRS-R decline^40^ and it is likely that there are several different patterns of decline in other measures of disease too. In contrast, our simple approach only relates the mean of one group to the mean of another. However, this approach can provide a starting point for more sophisticated attempts to reconcile preclinical and clinical disease measurement.

We suggest that translational biomarkers can have a key role to play in the design of preclinical studies. Clearly the use of “survival” in the SOD1^G93A^ model has been particularly unreliable as a predictor of clinical efficacy. By directly (or at least as directly as is possible), relating preclinical data to patient data using the same readout as we have done, preclinical studies can identify when the most clinically relevant starting point might be for the outcome measures being used. In addition, it may enable better predictions of treatment effect sizes for calculating clinical studies sample sizes. With further development, such approaches may be able to build confidence in the preclinical data package for successful translation of neuroprotective approaches into ALS clinical trials.

## Acknowledgements

This work was funded by Nxera Pharma UK Ltd.

## Author Contributions

S.M., A.K. and T.W. conducted the animal experiments. Z.Q and J.A. conducted the review and data extraction of the clinical studies. S.M., J.A. and R.M. prepared the manuscript.

## Competing Interests

The authors declare no competing interests.

## Data Availability Statement

The datasets analysed for human data are cited. Mouse data available on request.

## Notes

### Competing Interest Statement

The authors have declared no competing interest.

