## Supplemental Figure 1 for "Maximising the translational potential of neurophysiology in amyotrophic lateral sclerosis: a study on compound muscle action potentials"

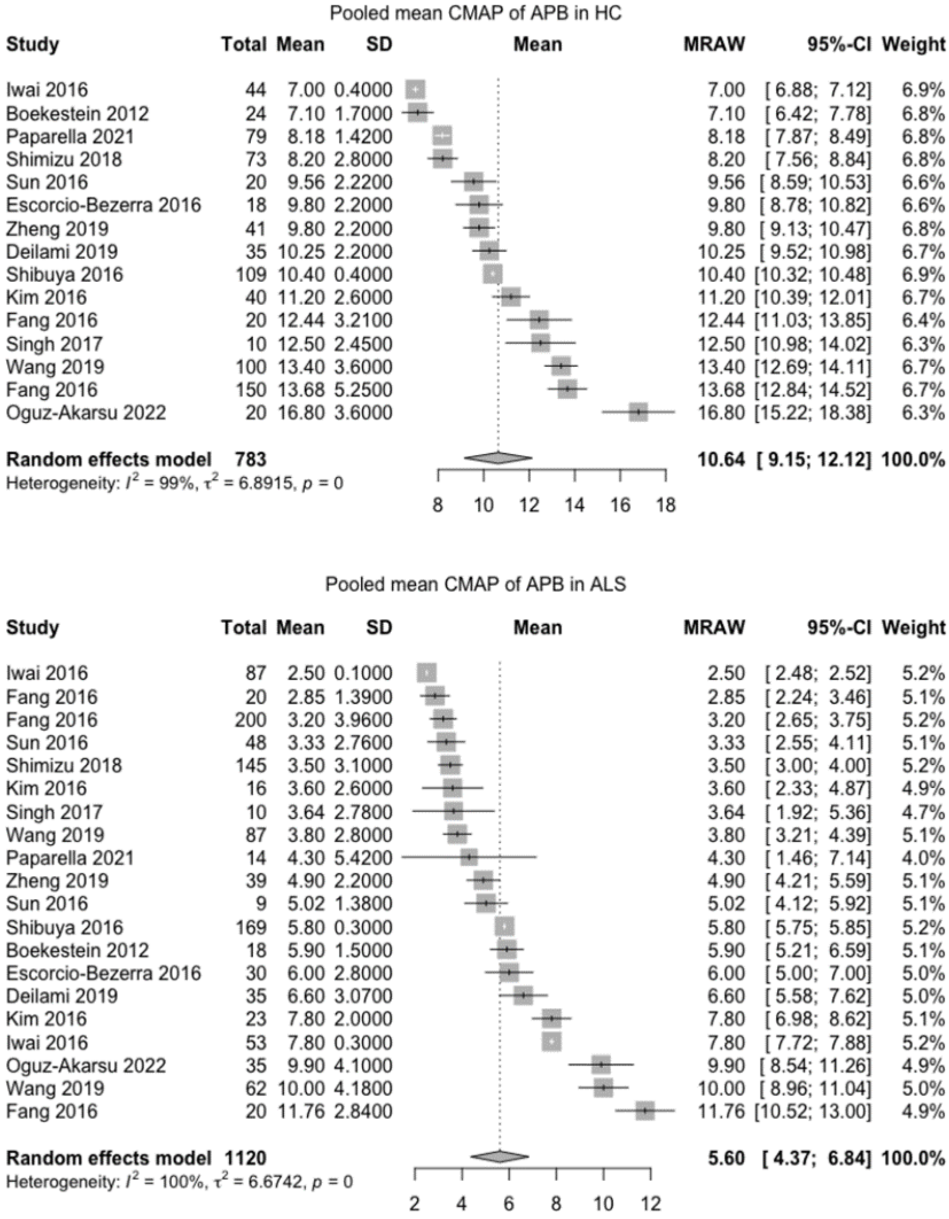


**Supplemental Figure 1. Study heterogeneity for APB.** Forest plots of pooled mean CMAP for healthy controls (HC, top) and ALS patients (ALS, bottom) derived from clinical studies.
