## Supplemental Figure 2 for "Maximising the translational potential of neurophysiology in amyotrophic lateral sclerosis: a study on compound muscle action potentials"

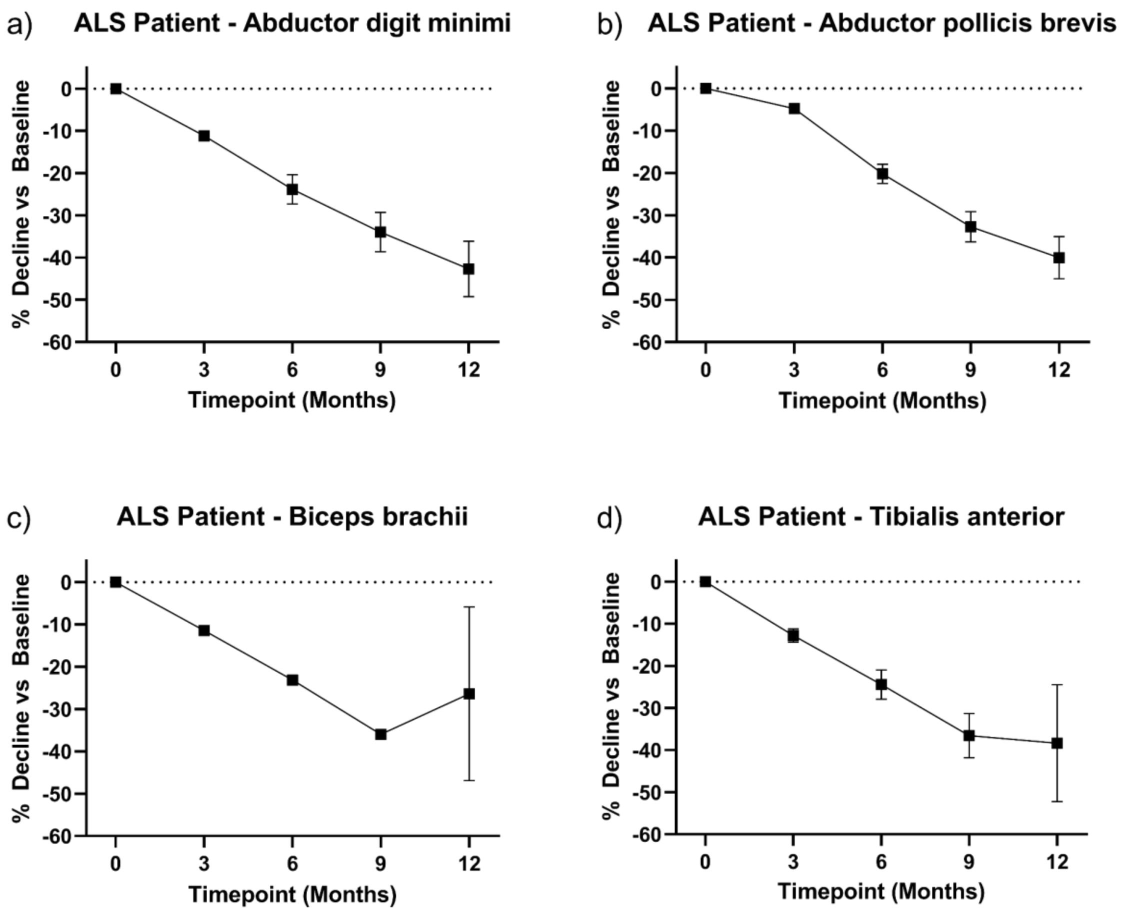


**Supplemental Figure 2. CMAP decline in ALS patients.** Average (± SEM) ALS patient CMAP percentage decrease versus baseline (first visit) for combined compound muscle action potential data gathered from multiple time point clinical studies in abductor digiti mini (a), abductor pollicis brevis (b), biceps brachii (c) and tibialis anterior (d).
