## Supplemental Figure 3 for "Maximising the translational potential of neurophysiology in amyotrophic lateral sclerosis: a study on compound muscle action potentials"

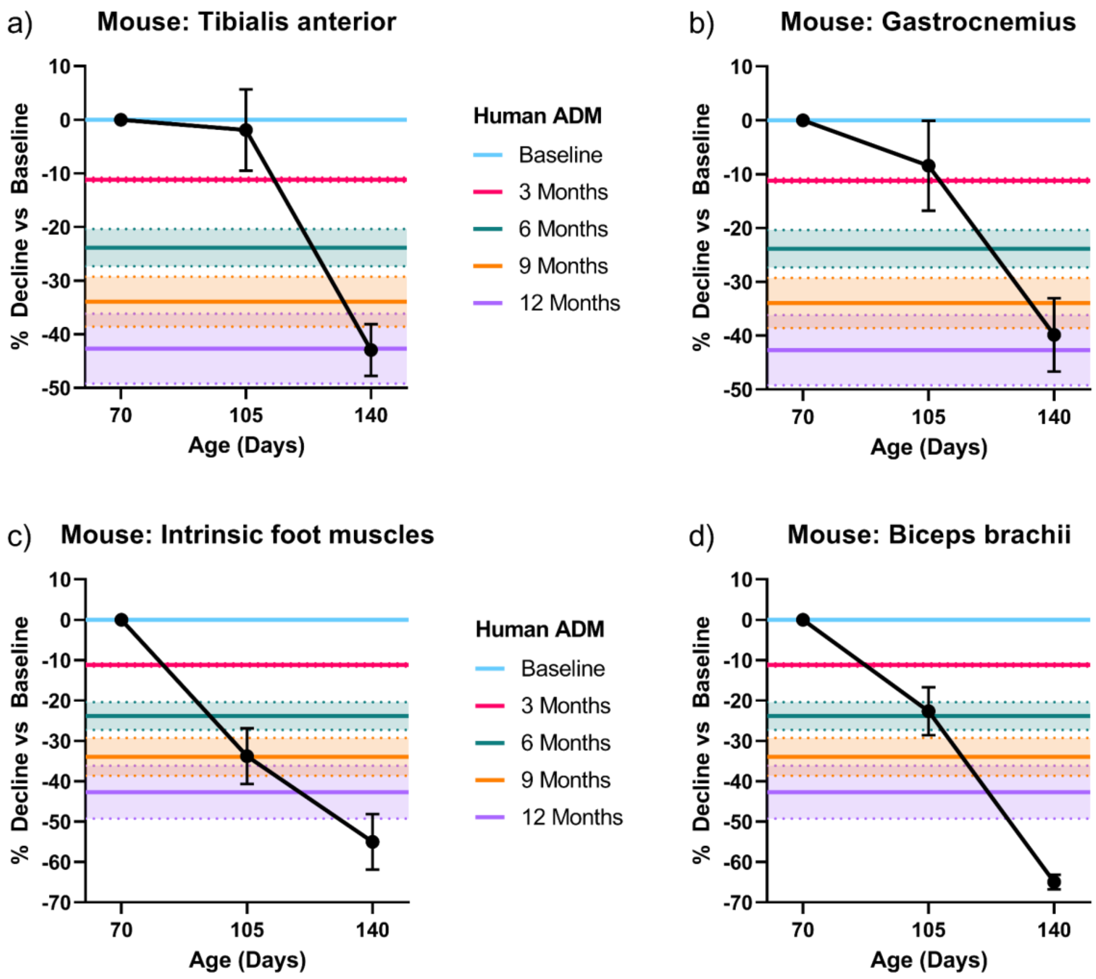


**Supplemental Figure 3. SOD1$^{G93A}$ tibialis anterior and gastrocnemius longitudinal data from P70-P140 represent a good match to the 12 month decline in human ADM.** Average (± SEM) CMAP decline compared to baseline recording at three time points in the mouse (black circles) in tibialis anterior (a), gastrocnemius (b), intrinsic foot muscle (c), and biceps brachii (d). The different time points in human ADM are represented by horizontal coloured lines (mean represented by the line and standard error by the shaded area).
