## Supplemental Figure 4 for "Maximising the translational potential of neurophysiology in amyotrophic lateral sclerosis: a study on compound muscle action potentials"

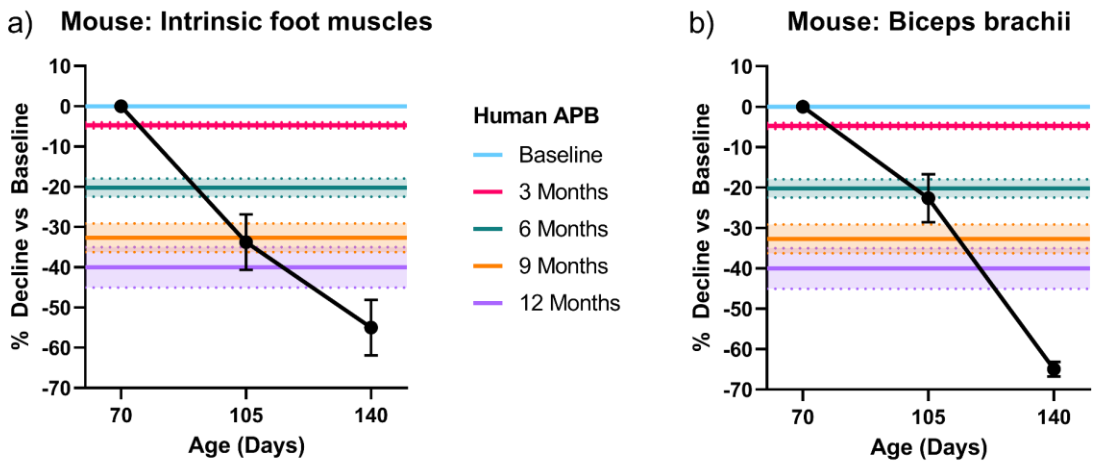


**Supplemental Figure 4. SOD1$^{G93A}$ intrinsic foot muscle and biceps brachii longitudinal data from P70-P105 most resembles the 6-9 month decline in human studies**. Average (± SEM) CMAP decline compared to baseline recording at three time points in the mouse (black circles) in the intrinsic foot muscle (a), and biceps brachii (b). The different time points in human APB are represented by horizontal coloured lines (mean represented by the line and standard error by the shaded area.
