## Supplemental Figure 5 for "Maximising the translational potential of neurophysiology in amyotrophic lateral sclerosis: a study on compound muscle action potentials"

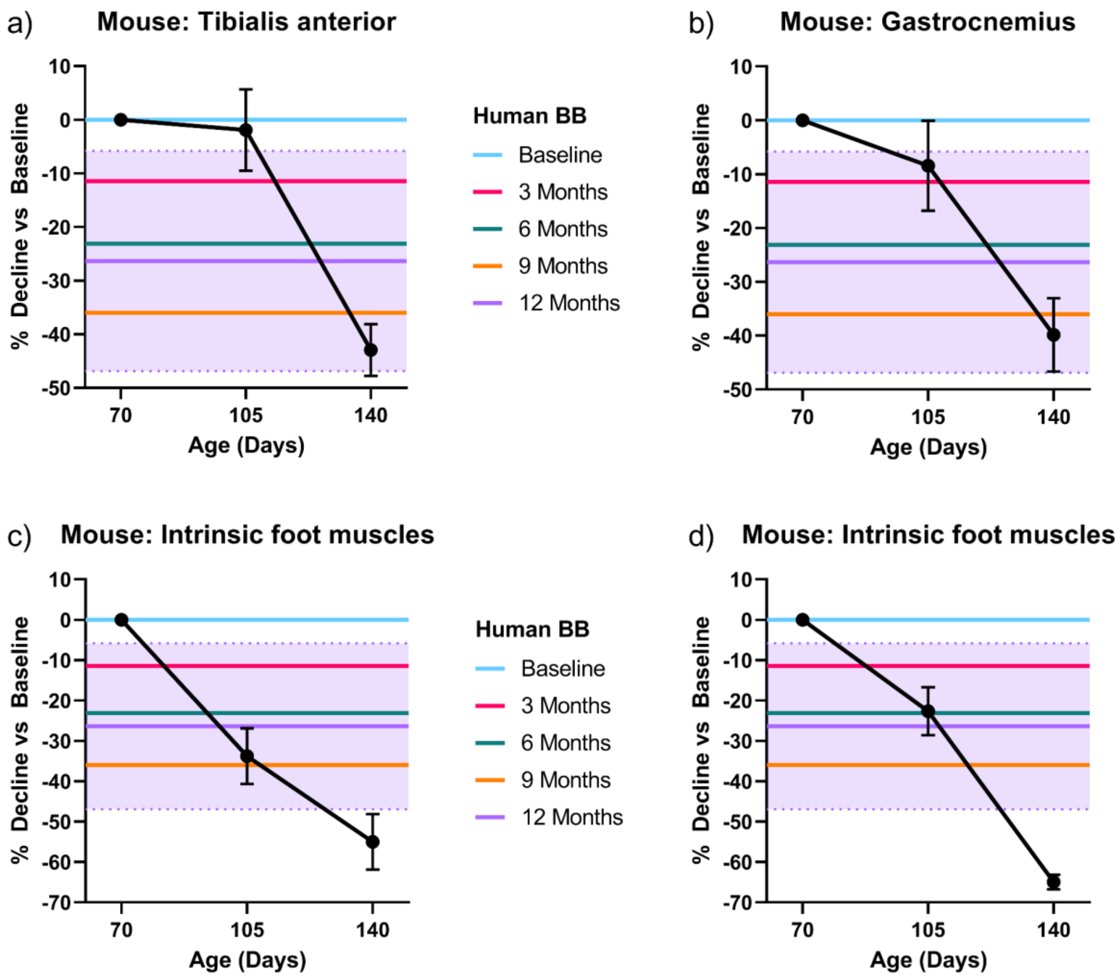


**Supplemental Figure 5. SOD1$^{G93A}$ tibialis anterior and gastrocnemius longitudinal data from P70-P140 most resembles the 9 month decline in human biceps brachii.** Average (± SEM) CMAP decline compared to baseline recording at three time points in the mouse (black circles) in tibialis anterior (a), gastrocnemius (b), intrinsic foot muscle (c), and biceps brachii (d). The different time points in human BB are represented by horizontal coloured lines (mean represented by the line and standard error by the shaded area).
