## Supplemental Table 1 for "Maximising the translational potential of neurophysiology in amyotrophic lateral sclerosis: a study on compound muscle action potentials"

**Supplemental Table 1. ALS patient CMAP percentage change against baseline summary data.** ALS patient CMAP percentage decrease (± SEM) versus baseline (first visit) for combined compound muscle action potential data gathered from multiple time point clinical studies. Note: the SEM for BB is small as only one study reporting the relevant data is retrieved from the literature.

| **Timepoint (Months)** | **Percentage Change Against Baseline (± SEM)** | | | |
|---|---|---|---|---|
| | 3 Months | 6 Months | 9 Months | 12 Months |
| **ADM** | -11.19% (± 0.38) | -23.85% (± 3.46) | -33.95% (± 4.68) | -42.70% (± 6.54) |
| **APB** | -4.77% (± 0.76) | -20.21% (± 2.29) | -32.73% (± 3.56) | -40.06% (± 5.01) |
| **BB** | -11.44% (± 0) | -23.14% (± 0) | -36.0% (± 0) | -26.37% (± 20.55) |
| **TA** | -12.79% (± 1.59) | -24.44% (± 3.49) | -36.56% (± 5.25) | -38.35% (± 13.87) |
