## Supplemental Table 1 for "Maximising the translational potential of neurophysiology in amyotrophic lateral sclerosis: a study on compound muscle action potentials"

**Supplemental Table 2. SOD1$^{G93A}$ CMAP percentage change against baseline summary data.** Percentage decrease (± SEM) versus baseline (70 days) in the SOD1G93A mouse compound muscle action potential data.

| **Time Point (Days)** | **Percentage Decline Against Baseline (± SEM)** | |
|---|---|---|
| | 105 days | 140 days |
| **Tibialis anterior** | -1.93% (± 7.58) | -42.93% (± 4.85) |
| **Gastrocnemius** | -8.42% (± 8.38) | -39.87% (± 6.82) |
| **Intrinsic Foot Muscles** | -33.80% (± 6.90) | -55.01% (± 6.89) |
| **Biceps brachii** | -22.67% (± 5.95) | -64.96% (± 1.83) |
